# Expression of Wilm’s Tumor Gene in Endometrium with Potential Link to Gestational Vascular Transformation

**DOI:** 10.1101/2020.03.11.987305

**Authors:** Peilin Zhang

**Author notes:** Correspondence: Peilin Zhang, MD., Ph.D., Pathology, New York – Presbyterian Brooklyn Methodist Hospital, 506 6^th^ St., Brooklyn, NY 11215.

## Abstract

**Background:** WT1 is a transcription factor with versatile cellular functions in embryonic development, maintenance of adult tissue functions and regenerations. WT1 is known to be regulated by progesterone and it is abundantly expressed in endometrium, but its function is unclear.

**Design:** WT1 expression was detected by immunohistochemical staining in endometrium of various physiological and pathological conditions.

**Result:** WT1 was detected in endometrial stromal cells and vascular smooth muscle cells in both proliferative and secretory phases of menstrual cycles. WT1 appeared increased in vascular smooth muscle cells in spiral artery in early pregnancy and WT1 was also detected in regenerative endothelial cells and smooth muscle cells in decidual vasculopathy at term. WT1 expression was decreased in endometrial stromal cells in adenomyosis (endometriosis).

**Conclusion:** WT1 potentially links the hormonal (progesterone) effects on endometrial decidualization and may play a role in gestational vascular transformation during pregnancy and restoration after pregnancy.

## Introduction

Endometrial decidualization is characterized by morphological transformation of epithelium, endometrial stromal tissue and terminal segments of the spiral artery through the action of steroid hormone, progesterone, secreted by the corpus luteum after ovulation [1, 2]. Embryonic implantation occurs in a narrow window of time after fertilization and the embryo attaches and invades into endometrial tissue through complex interactions between the developing trophectoderm and the decidualized endometrium [2]. Extravillous trophoblasts derived from the early trophectoderm invade into the decidualized endometrial stromal tissue and the spiral artery, leading to trophoblasts-dependent spiral artery remodeling by forming the endovascular trophoblastic plugs with concurrent replacement of smooth muscle wall and endothelium [3]. Uterine natural killer cells (NK) play critical roles in trophoblasts dependent spiral artery remodeling and the formation of endovascular trophoblastic plugs is associated with phenotypic switch of the endovascular trophoblasts to express CD56, a defining molecular marker for NK cell lineage [4–7]. Meanwhile, endometrial spiral artery is undergoing trophoblasts independent remodeling morphologically characterized by arterial wall hypertrophy/hyperplasia (mural hypertrophy/hyperplasia) and endothelial vacuolation, and this morphological transformation occurs distant from the extravillous trophoblasts without direct trophoblastic interaction [8]. The molecular mechanism of trophoblasts independent remodeling of spiral artery (mural hypertrophy/hyperplasia) is likely through endocrine (hormonal) actions but the precise hormonal action on the smooth muscle cells of the vascular wall is poorly understood. In endometrial decidualization and subsequent embryonic implantation, progesterone and human chorionic gonadotropin (hCG) actions are predominant for decidual, trophoblastic and fetal development, and the maintenance of pregnancy in first trimester, and the critical role of these hormones on spiral artery remodeling is yet to be defined [2].

Wilm tumor 1 gene (WT1) is a tumor suppressor gene discovered in patients of Wilm’s tumor, a pediatric malignancy with versatile molecular functions [9–14]. WT1 is a transcription factor regulating a large number of target gene expressions mediating a variety of cellular functions in organ development, homeostasis and diseases [15, 16]. Mutations of WT1 gene were shown to be associated with Denys-Drash syndrome and Frasier syndrome, and deletion of the short arm of the chromosome 11 containing WT1 gene is associated with WAGR syndrome (Wilm’s tumor, anirida, genitourinary anomalies, and mental retardation) [14, 17, 18]. WT1 is interacting with many different partner proteins important for various cellular activities [9, 14]. Normal WT1 gene function is critical for development of urogenital system, mesothelial cell function and vascular endothelial cell functions [14]. Clinically WT1 is also important for adult kidney function and cardiovascular functions [14, 19]. In pathology practice, immunohistochemical staining for WT1 serves as a marker for ovarian carcinoma, mesothelial cell and endothelial cell lineages [20, 21]. There is abundant WT1 gene expression in the endometrium by RNA microarray studies (Human Protein Atlas https://www.proteinatlas.org/), but the function of WT1 in endometrium is yet to be defined, especially when the endometrium undergoes significant changes during menstrual cycle and the pregnancy. As initial steps, expression of WT1 protein in endometrium by immunohistochemical staining is attempted to locate the WT1 protein during menstrual cycles and pregnancy.

## Material and methods

The study is exempt from Institutional Review Board (IRB) approval according to section 46.101(b) of 45CFR 46 which states that research involving the study of existing pathological and diagnostic specimens in such a manner that subjects cannot be identified is exempt from the Department of Health and Human Services Protection of Human Research Subjects. Endometrial and placental tissues submitted for pathology examination for a variety of clinical indications were included in the study using the routine Hematoxylin & Eosin (H&E) stain. Proliferative endometrium, secretory endometrium, decidua and implantation sites were obtained from the submitted pathology specimens during clinical practice. Second trimester placentas were obtained from inevitable abortion specimens due to chorioamnionitis. Paraffin-embedded tissues from routine surgical pathology specimens and routine H&E stained pathology slides were examined by light microscopy using the Amsterdam criteria for placental examination [22]. Immunohistochemical staining for WT1 was used to highlight the specific cell types of the muscular vessels, endometrium and myometrium, fetal villous stem villous arteries and umbilical cord vessels. CD34 was also used to highlight the endothelium of the endometrial or decidual vessels in comparison. Immunohistochemical staining procedures were performed on paraffin-embedded tissues using Leica Biosystems Bond III automated immunostaining system following the manufacturer’s instructions. WT1 and CD34 monoclonal antibodies were purchased for clinical applications from Dako Agilent under the catalogue number M3561 and M7165.

Totally 3 cases of proliferative endometrium, 3 cases of secretory endometrium, 2 cases of therapeutic abortion specimens at 10 weeks gestation, 1 case of inevitable abortion at 17 weeks, 4 cases of term placentas and umbilical cords at 39-40 weeks with and without decidual vasculopathy were examined by light microscopy and H&E stain as well as immunostaining for WT1 and CD34.

## Results

In small muscular arteries or arterioles of the lower extremities and the ovaries, WT1 was detected by immunohistochemical staining within the endothelial cells only. The staining signals appeared to be both cytoplasmic and nuclear. The smooth muscle layers of the vascular walls showed no or minimal immunostaining signals (Figure 1). The adventitia layer of the artery was evident on H & E staining slides, but no WT1 signals were present. In proliferative endometrium, WT1 expression was present in the nuclei of stromal cells and the endothelial cells as well as the smooth muscle cells of the spiral artery (Figure 2, top panel). The endometrial glands and epithelium and lymphocytes showed no staining signals. In the myometrium, the endothelial cells of the artery showed WT1 staining signals as well as the entire myometrium but no smooth muscle cells of muscular artery were staining for WT1 (Figure 2 bottom panel). The adventitia layer was not evident in the endometrial arteries, and no convincing staining signals for WT1 were noted for the adventitial layers of myometrial arteries. In contrast to those of the proliferative endometrium, the endometrial stromal cells in adenomyosis showed no nuclear staining signals for WT1 (Figure 3 bottom panel). In secretory endometrium, WT1 expression was identified in the stromal cells, endothelial cells and smooth muscle cells of the spiral artery walls (Figure 4 top panel), similar to those of proliferative endometrium (Figure 2). There appeared to be stronger staining signals for WT1 in smooth muscle layers of spiral artery but the immunohistochemical staining method was not calibrated for quantitative measurement of WT1 expression. WT1 was seen in the endothelial cells of the myometrial segment of the spiral artery, and no significant WT1 signals were present in the smooth muscle layers or adventitial layers (Figure 4, bottom panel).

**Figure 1:**
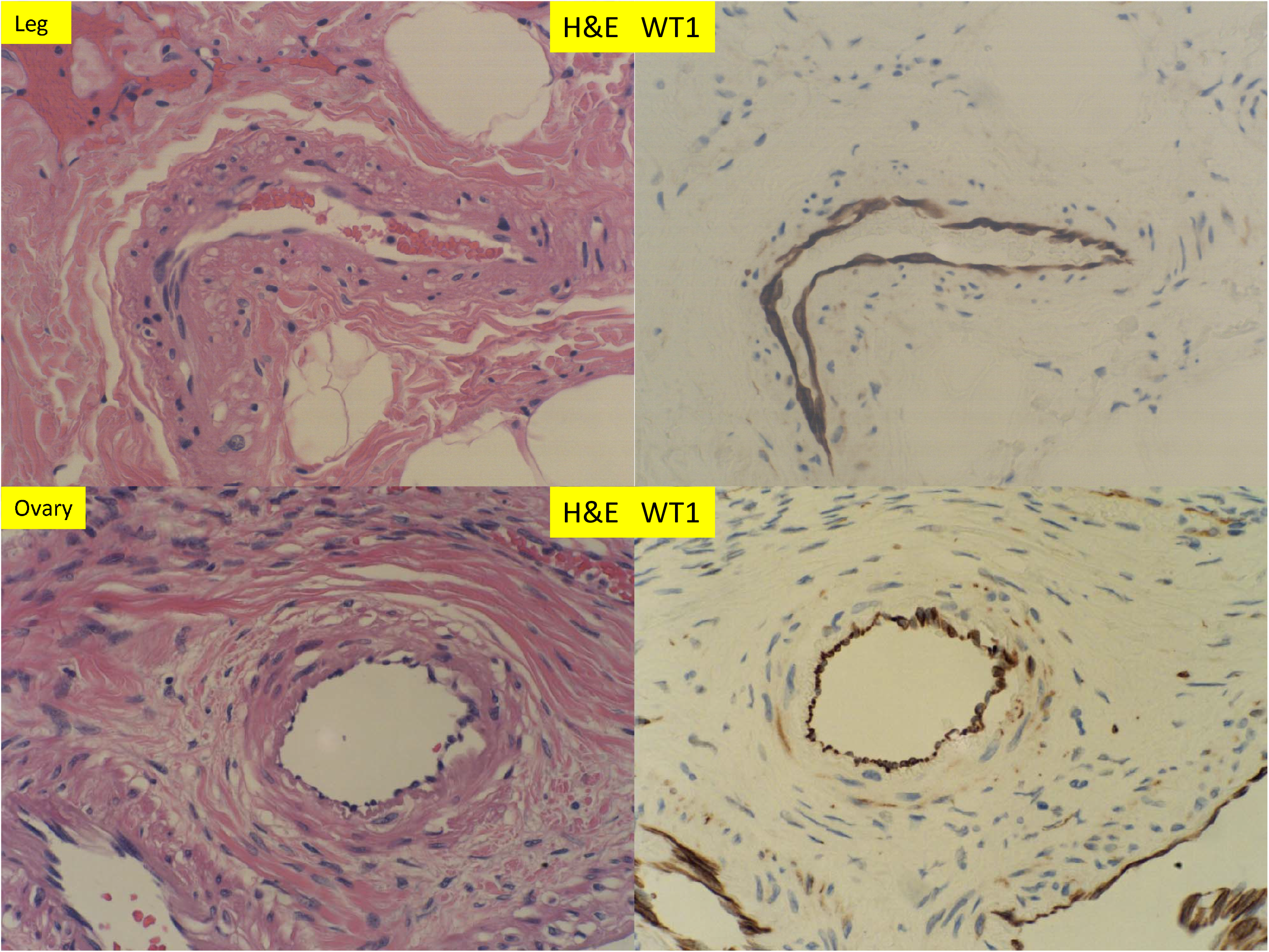
H & E staining and immunostaining for WT1 expression of muscular arteries in the soft tissue of the lower leg after traumatic amputation and the ovary tissue of peri-menopausal women (400 x magnification).

**Figure 2:**
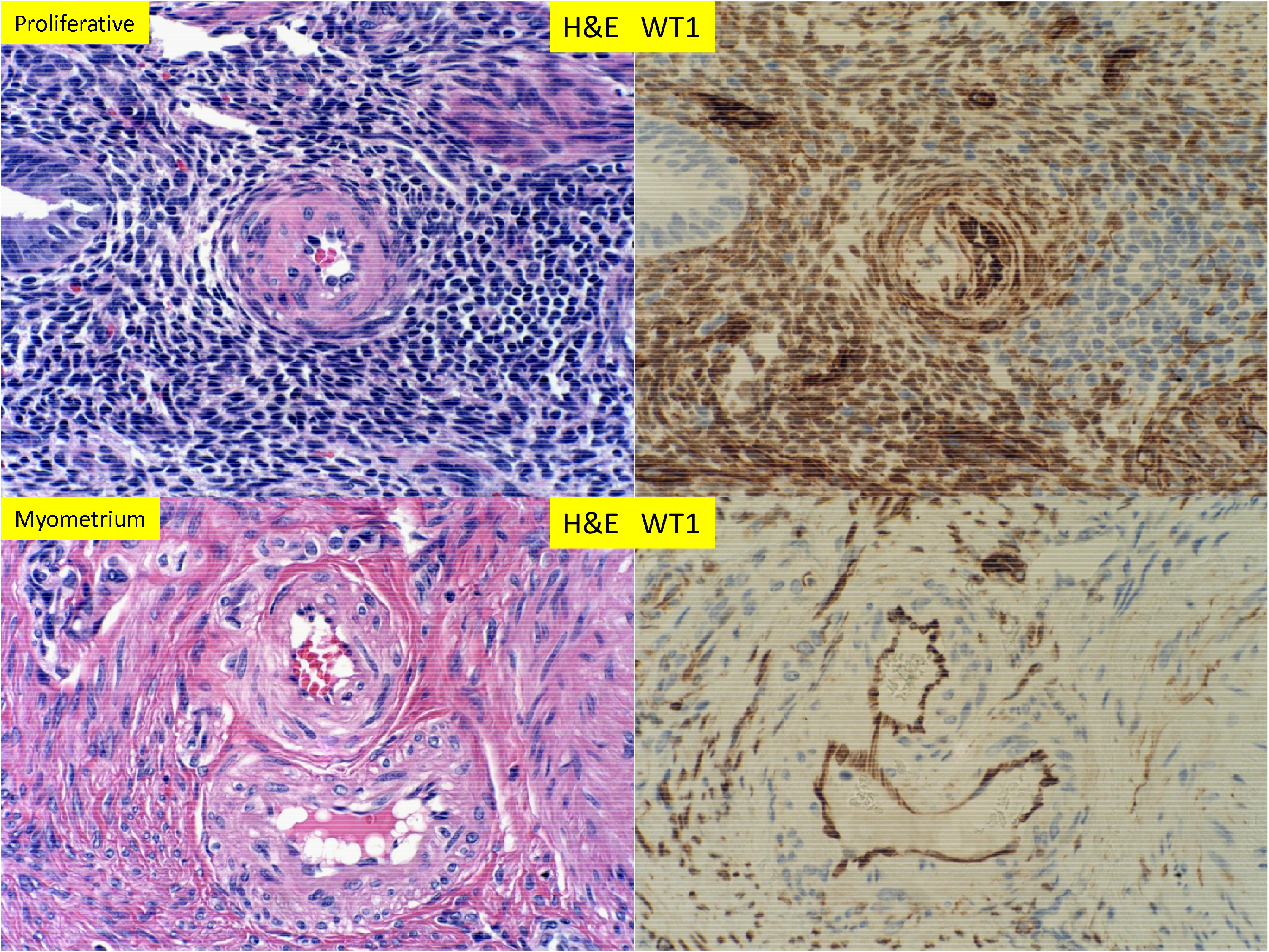
H & E staining and immunostaining for WT1 expression of proliferative endometrium with glandular, stromal cells and spiral artery and myometrium with spiral artery (400 x magnification).

**Figure 3:**
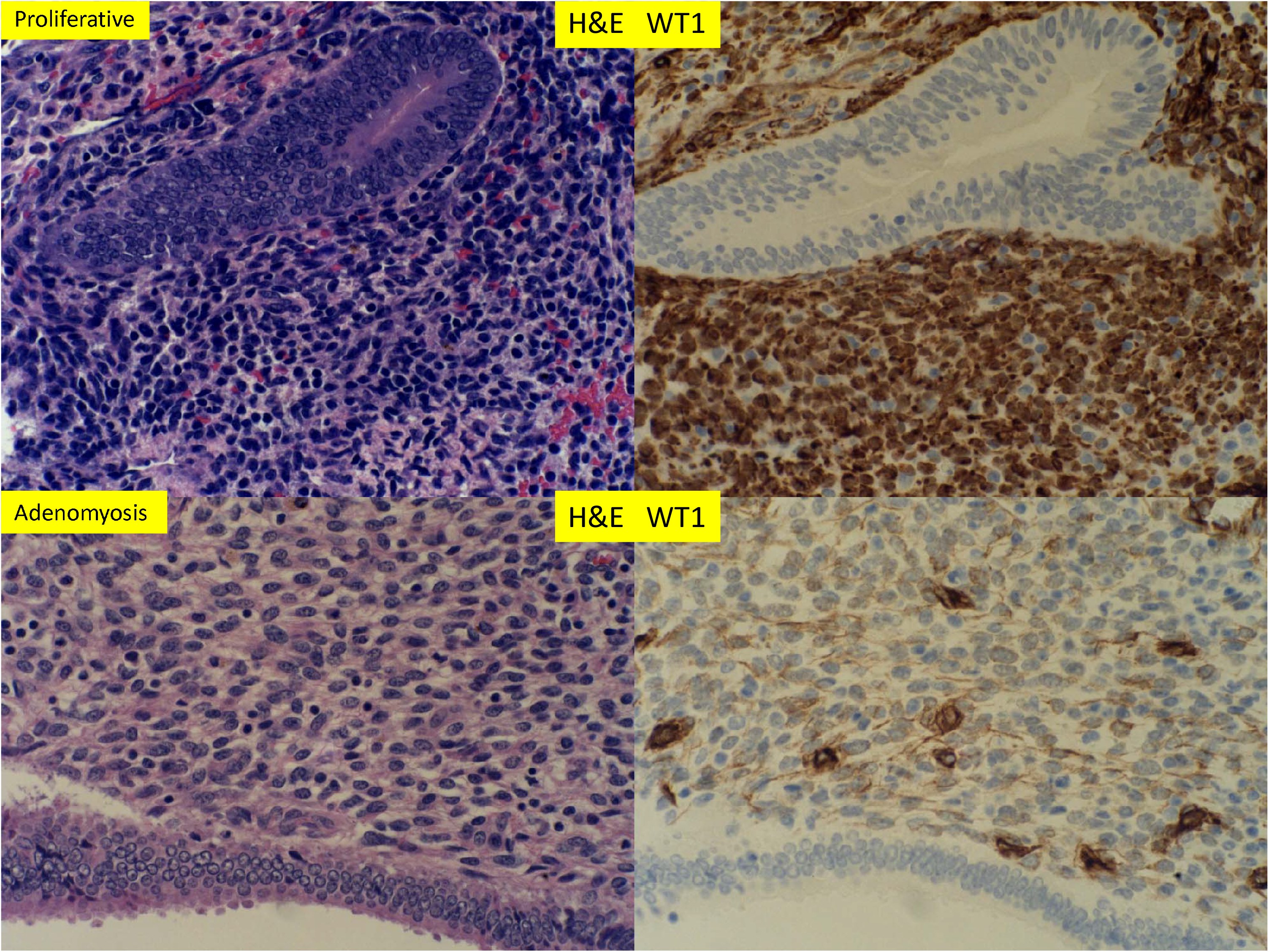
H & E staining and immunostaining for WT1 expression of proliferative endometrium and adenomyosis in hysterectomy specimen. Noted a significant reduction of WT1 signals within the stromal cells (400 x magnification).

**Figure 4:**
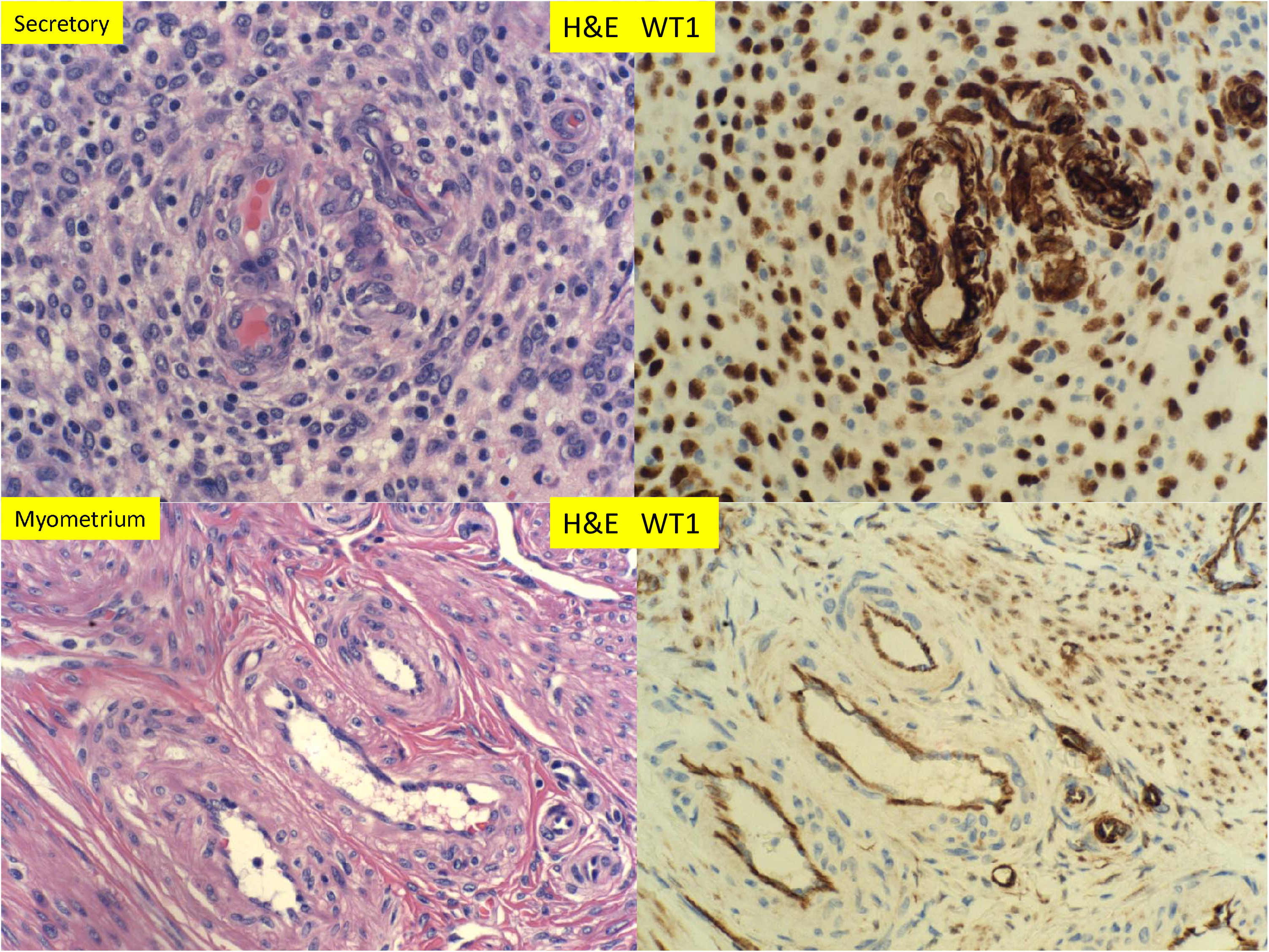
H & E staining and immunostaining for WT1 expression of secretory endometrium and myometrium (400 x magnification)

In early pregnancy at 10 weeks when the spiral arteries underwent remodeling in both the trophoblasts dependent and independent manners, the decidual cells expressed WT1 strongly within the nuclei (Figure 5). There were strong nuclear signals for WT1 in hyperplasia/hypertrophy of the smooth muscle walls of the spiral artery (Figure 5, top panel). There appeared to be cytoplasmic WT1 signals within the endothelial cells in the arterial lumen. In contrast, the spiral arteries with trophoblastic plugs and the complete replacement of muscular walls were negative for WT1 expression (Figure 5, bottom panel). At 17 weeks gestation (Figure 6), the expression patterns of WT1 in decidua, spiral artery with hypertrophy/hyperplasia and those with trophoblastic plugs were similar to those at 10 week gestations (Figure 5). There was no difference in WT1 expression pattern and intensity between those of 10 weeks and 17 week gestation. No WT1 staining signals were noted in the adventitia layers of vessels at either 10 weeks or 17 weeks gestation.

**Figure 5:**
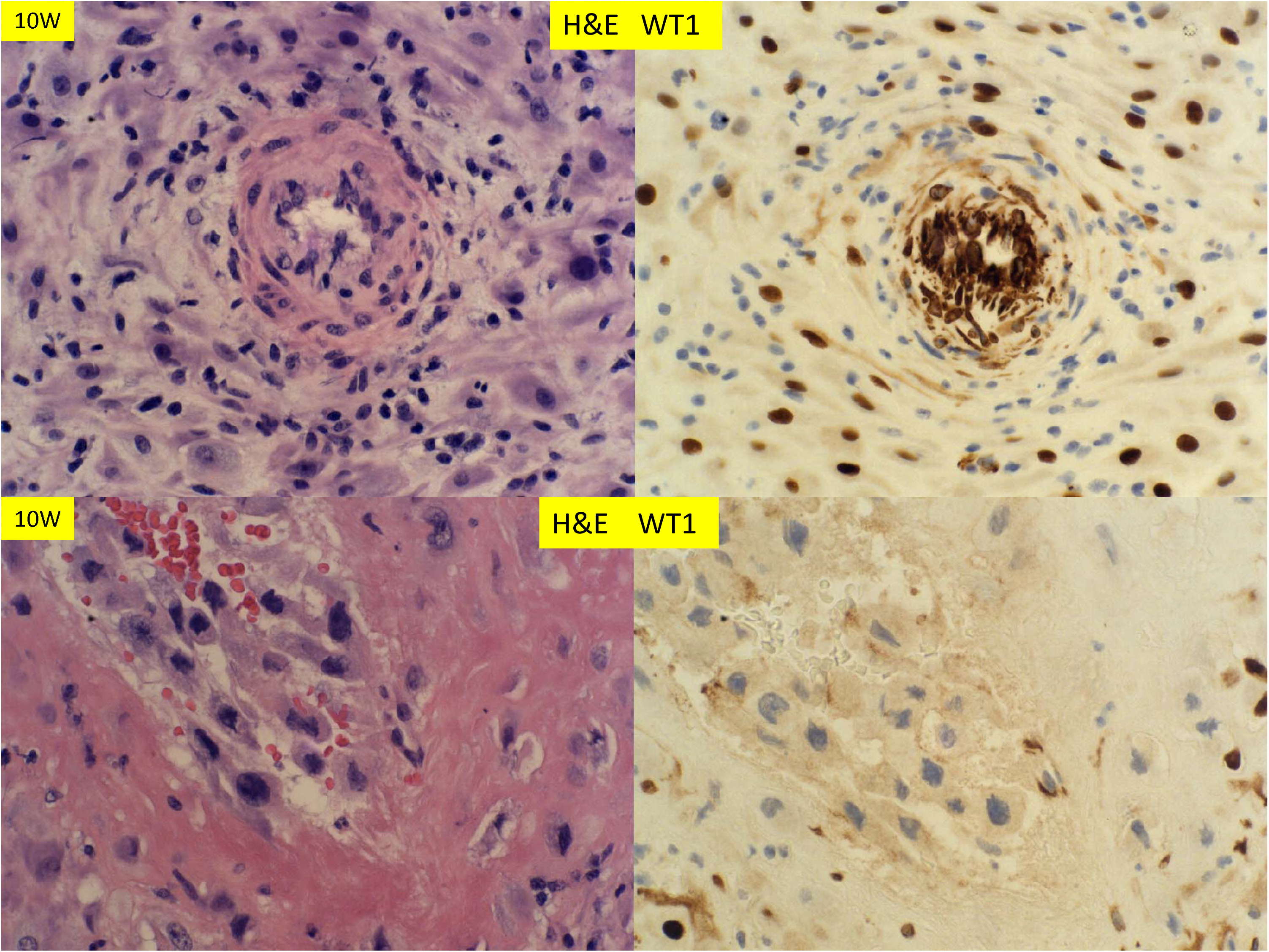
H & E staining and immunostaining for WT1 expression of decidua at 10 week gestation of therapeutic abortion. Spiral artery with mural hypertrophy/hyperplasia (top panel) and endovascular trophoblastic plug (bottom panel) (400 x magnification)

**Figure 6:**
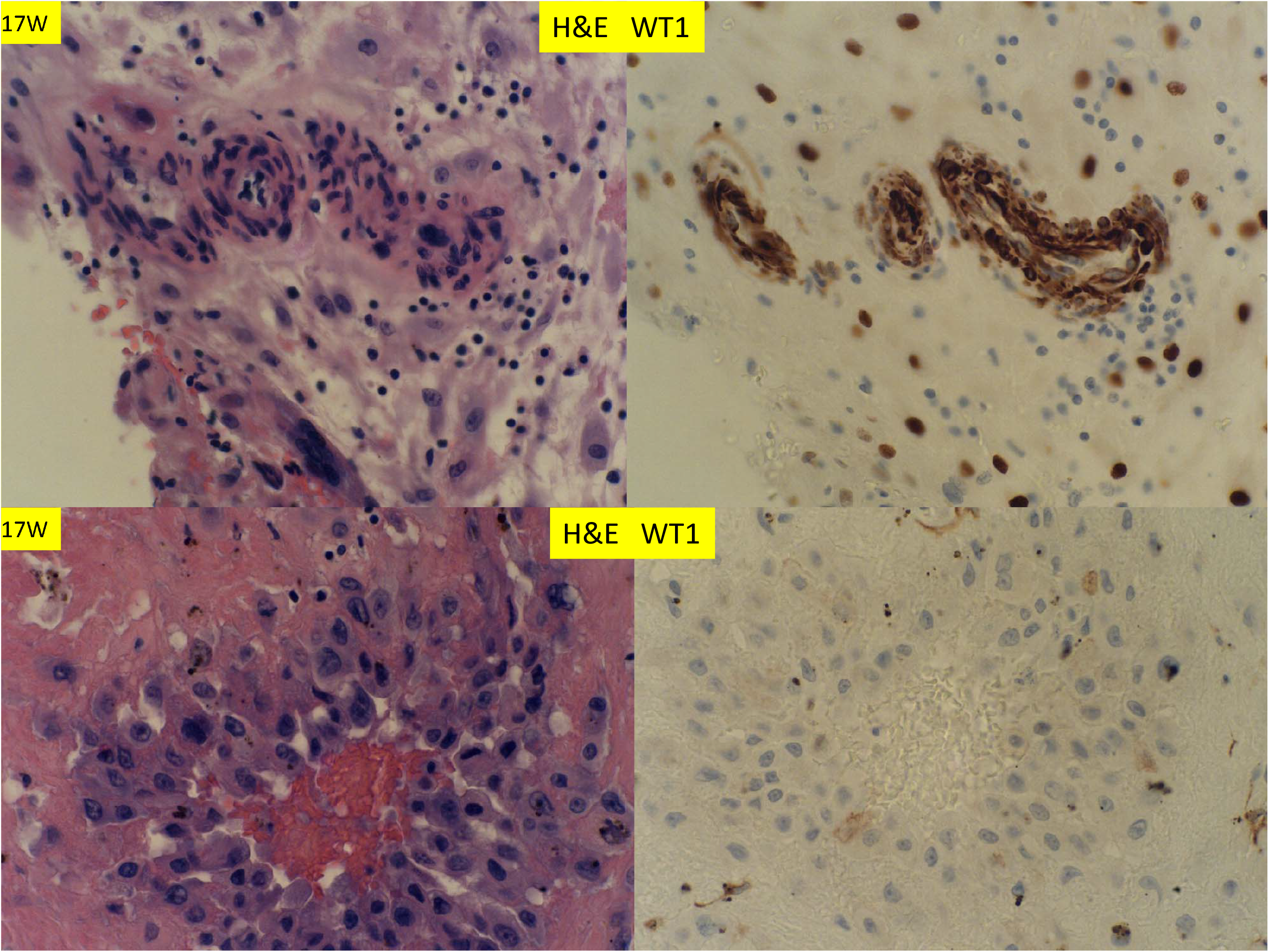
H & E staining and immunostaining for WT1 expression of decidua at 17 weeks gestations with inevitable abortion and chorioamnionitis. Spiral artery with mural hypertrophy/hyperplasia (top panel) and endovascular trophoblastic plug (bottom panel) (400 x magnification)

There are two types of decidual vasculopathy, mural arterial hypetrophy/hyperplasia and classic type including atherosis and fibrinoid medial necrosis. Mural hypertrophy/hyperplasia in the membrane roll (decidual vera) showed WT1 expression in endothelial cells and smooth muscle cells (Figure 7 top panel), and it appeared that the expression in the smooth muscle cells was decreased in comparison to those in early pregnancy (Figure 5 and 6). There was no WT1 expression in classic decidual vasculopathy including atherosis and fibrinoid medial necrosis. However, WT1 was detected in the regenerating endothelial cells and smooth muscle walls (Figure 7 bottom panel), and fibrinoid medial necrosis was partially present and negative for WT1 expression in the vessel with regenerating endothelium and smooth muscle wall (Figure 7 bottom panel). To illustrate this point of regeneration further, endovascular trophoblasts can be identified by immunostaining for CD56, and regenerating smooth muscle cells for smooth muscle myosin heavy chain (SMYOHC), as well as WT1 expression (Figure 8).

**Figure 7:**
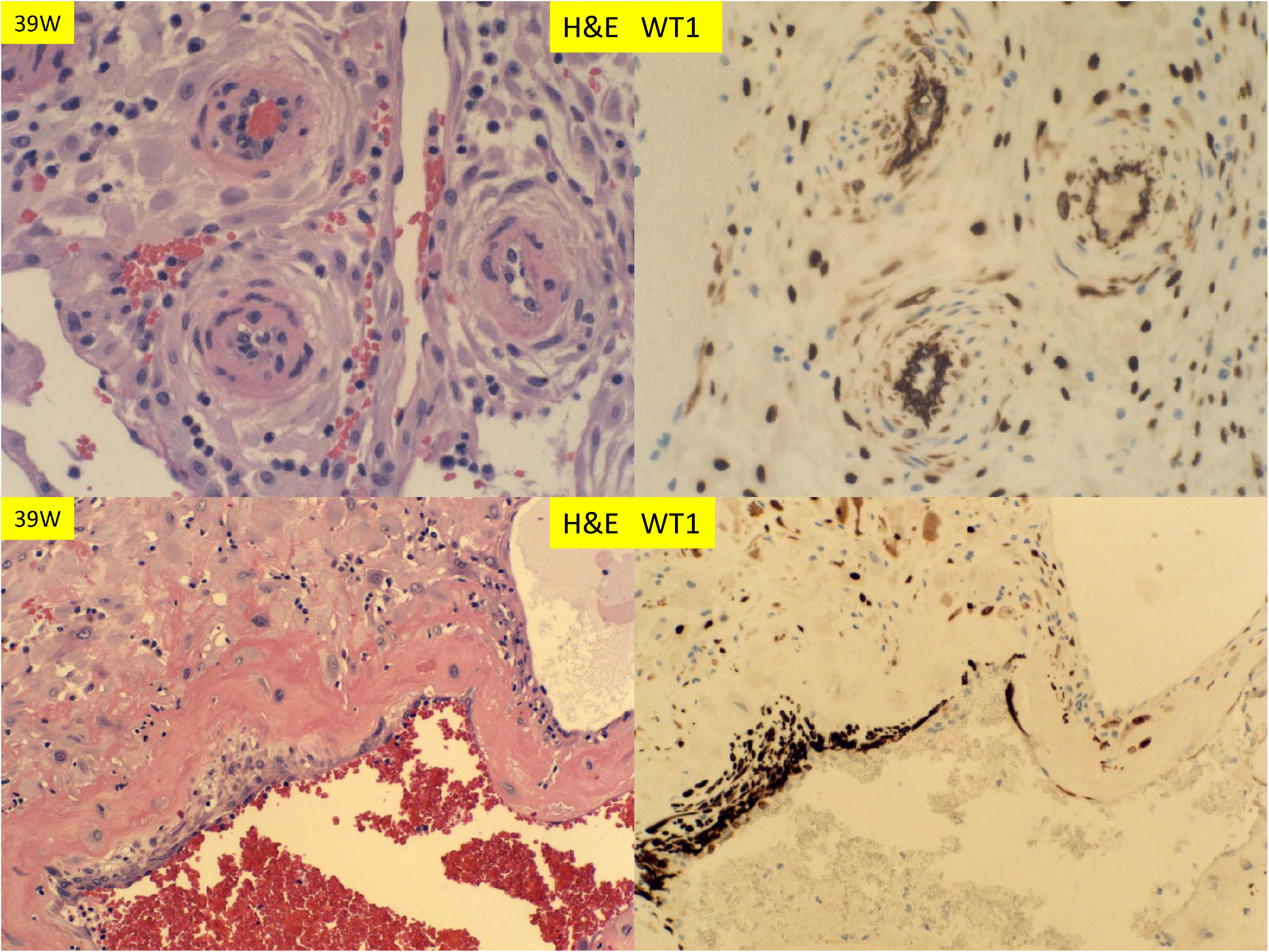
H & E staining and immunostaining for WT1 expression of decidual vessels in the membrane roll (vera) and decidua basalis at 39 weeks (decidual vasculopathy). Spiral artery at decidua vera (top panel) and partially restoring spiral artery after vasculopathy (bottom panel) (400 x magnification)

**Figure 8:**
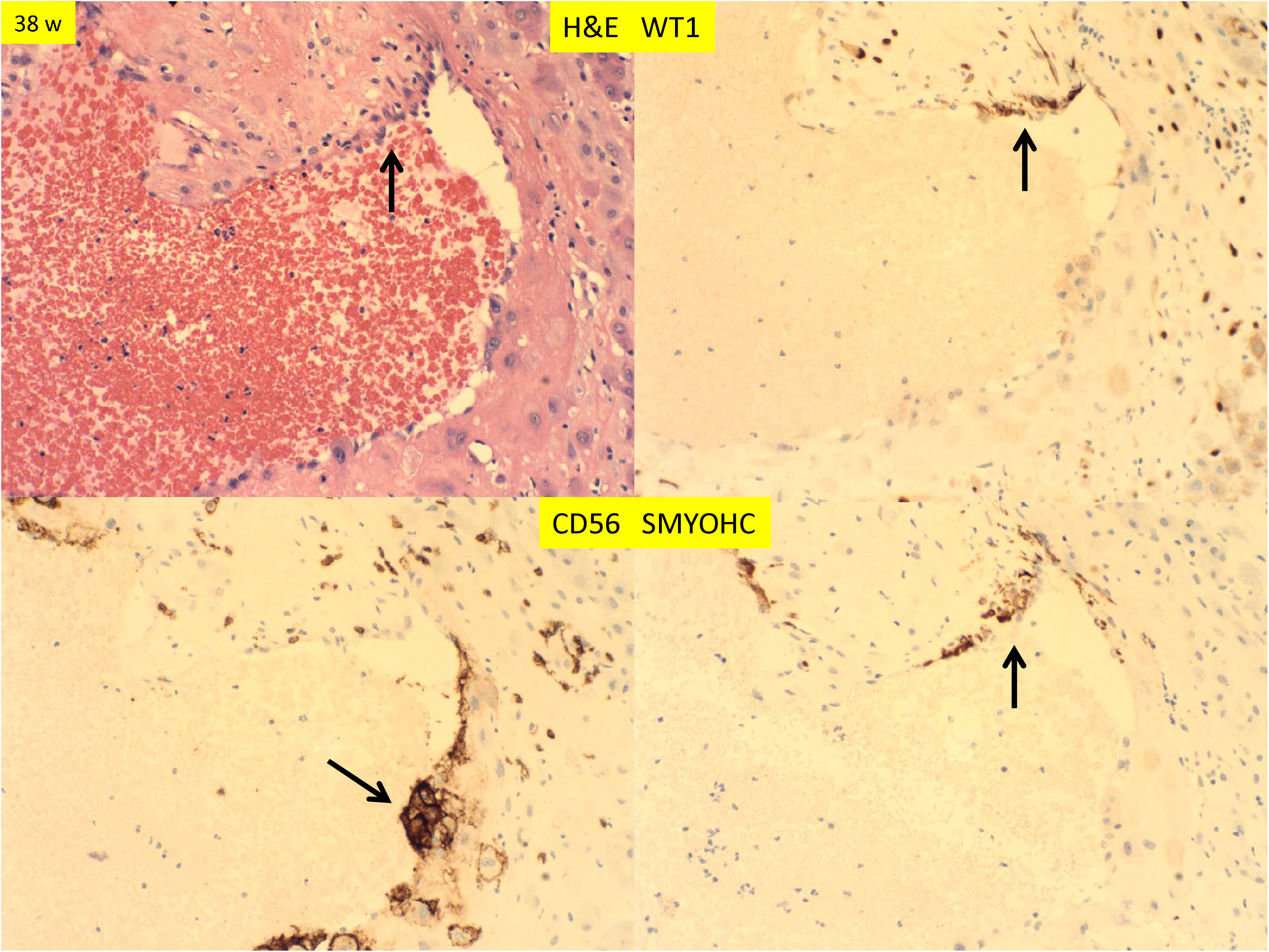
H & E staining and immunostaining for WT1, CD56 and smooth muscle myosin heavy chain (SMYOHC) expression of decidual vessels in the membrane roll (vera) and decidua basalis at term (decidual vasculopathy) at 38 weeks. Arrows indicate the focus of regenerating endothelium and smooth muscle cells with the remaining endovascular trophoblasts (200 x magnification)

In tubal pregnancy, WT1 expression was noted to be abundant in the tubal epithelium and smooth muscle wall and endothelium of the vessels, as well as small capillary endothelial cells (Figure 9). This is in contrast to the endometrium where WT1 was only expressed in the endometrial stromal cells, not in the endometrial glandular epithelium.

**Figure 9:**
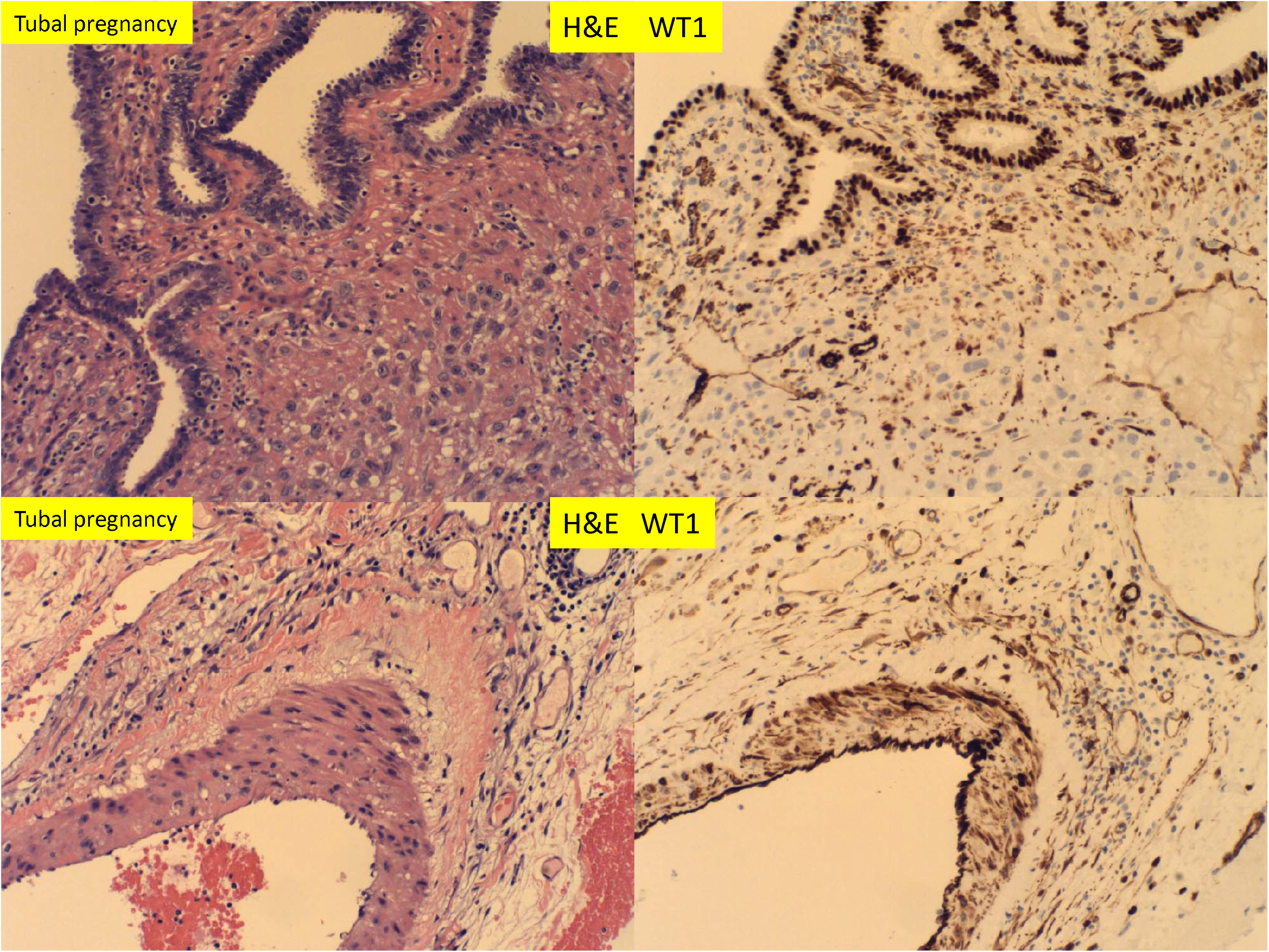
H & E staining and immunostaining for WT1 expressions of tubal pregnancy at 11 weeks after salpingectomy. Noted the tubal epithelium and tubal vessels with WT1 expression (400 x magnification)

In placental stem villi at term, the muscular artery showed intense WT1 staining signals but the luminal endothelial cells appeared no staining (Figure 10 top panel). This WT1 staining pattern in stem villous artery was in direct contrast to CD34 in which only the luminal endothelial cells showed CD34 staining signals (Figure 10 top and bottom panels). The WT1 expression patterns in stem villous artery were similar to those in the umbilical artery and veins in umbilical cord at term (Figure 11). There were no adventitia layers of these vessels as these vessels are anatomically different from those in the systemic circulation [23].

**Figure 10:**
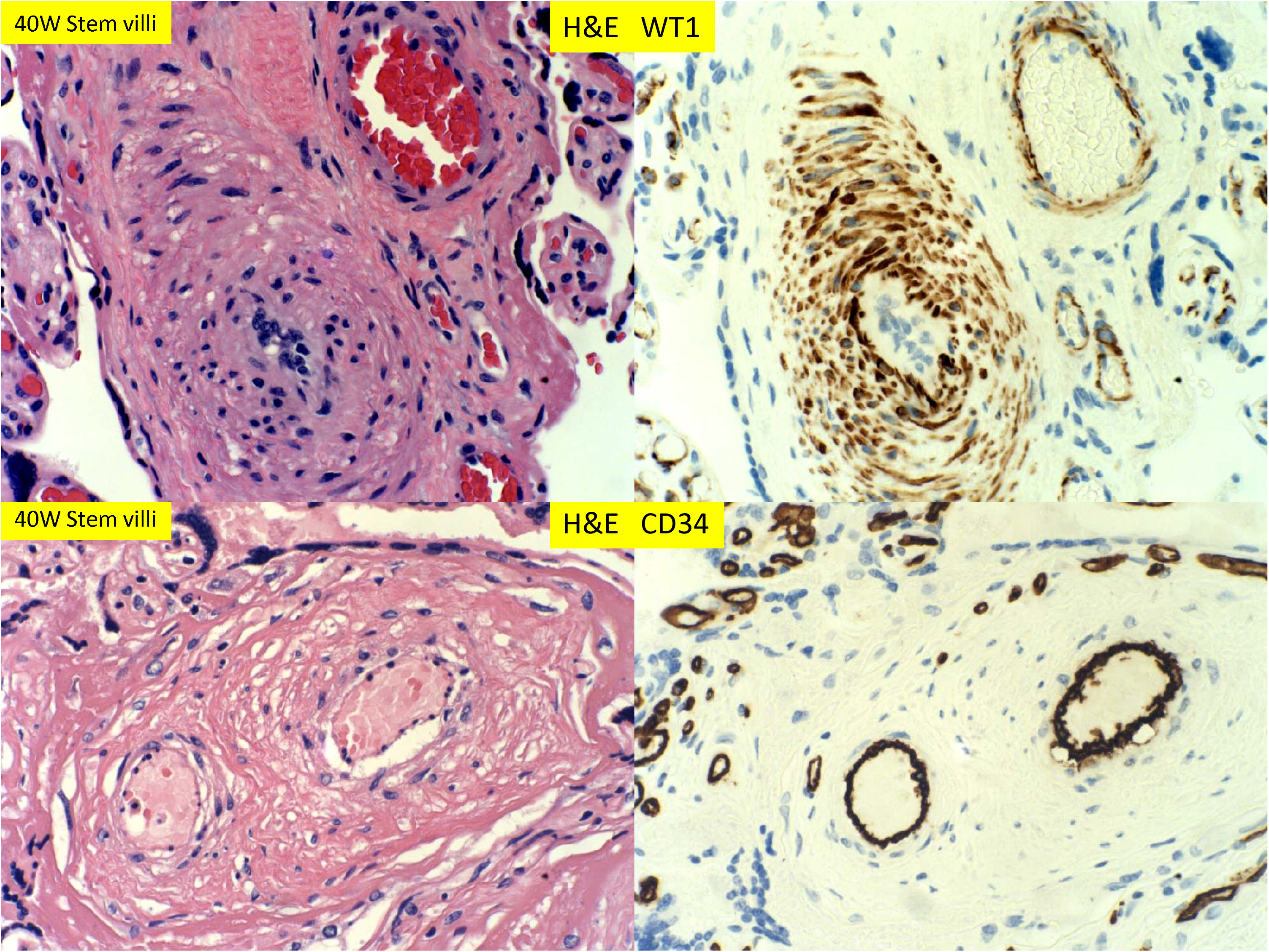
H & E staining and immunostaining for WT1 and CD34 expression of muscular arteries of stem villi of 39 week placentas. WT1 was noted in smooth muscle wall but not in endothelium whereas CD34 was only seen in endothelium (400 x magnification).

**Figure 11:**
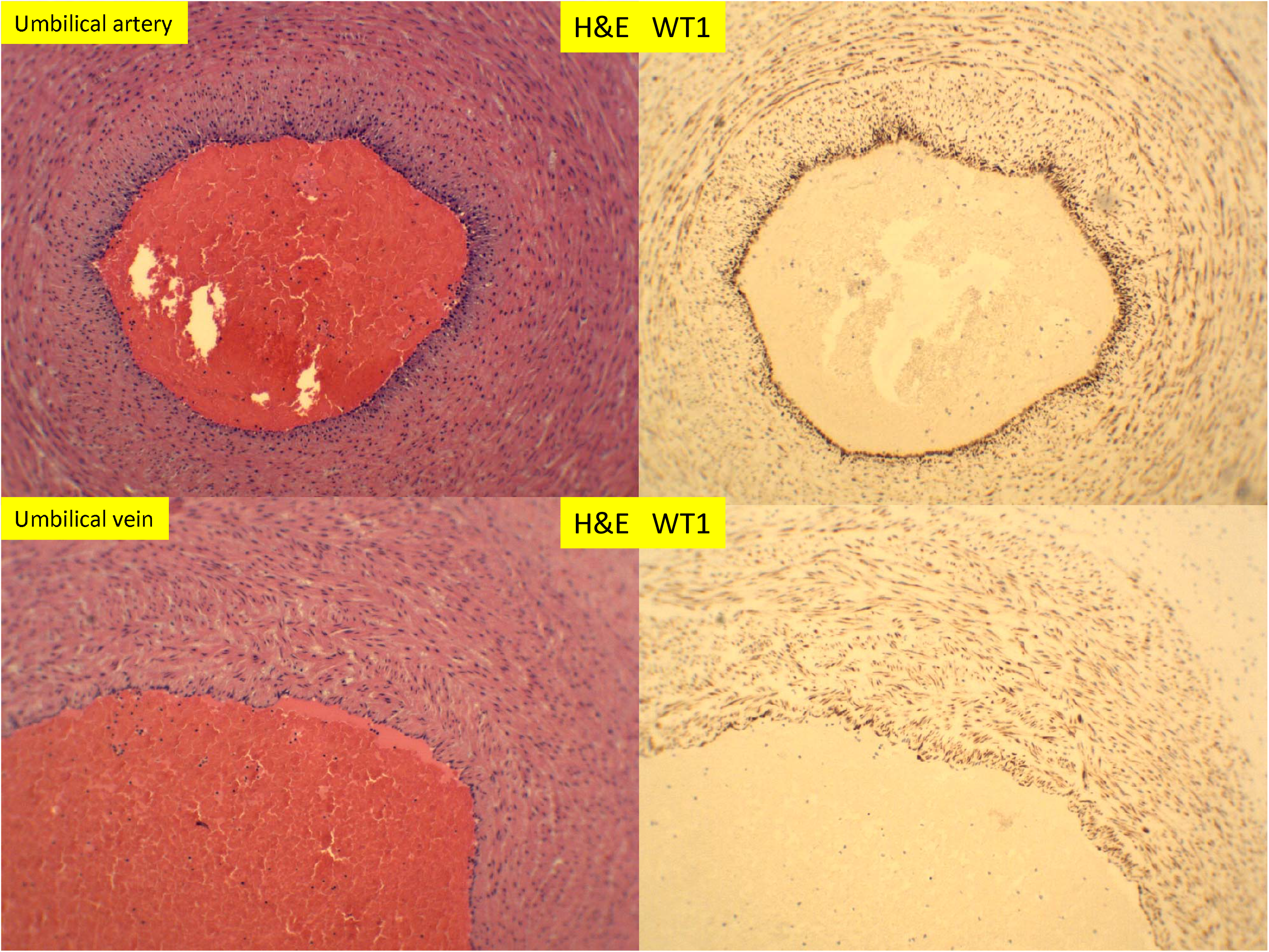
H & E staining and immunostaining for WT1 expression of umbilical artery and vein of 39 week placenta (100 x magnification)

## Discussion

WT1 gene is important for organogenesis of urogenital system, endothelial cell system and mesothelial cell system during embryogenesis, and its versatile cellular function is likely to be conferred through not only regulating multiple target gene expression but also receiving and mediating effects of multiple factors in growth and development [9, 14]. The WT1 function in endometrium appeared at least in part to be through regulation by the progesterone hormone and insulin – like growth factors (IGF), both of which are known to be critical for decidualization, implantation and fetal development [24–26]. WT1 has been shown to be up-regulated by progesterone in in vitro models, and it can also be self-regulated [24, 27]. Our current WT1 expression data in endometrium is consistent with the known function of progesterone in both menstrual cycles and pregnancy. However, the basal expression of WT1 in the stromal cells of proliferative endometrium and the epithelium of the fallopian tubes appear not related to the progesterone effect, and there is no estrogen or progesterone receptor expression in the tubal epithelium (data not shown). In current study, the WT1 expression is abundant in the stromal cells of the secretory endometrium and the decidua in first and second trimesters as well as the smooth muscle wall of the spiral arteries. The spiral artery transformation in secretory phase of endometrium and remodeling in early pregnancy characterized by mural arterial hypertrophy/hyperplasia correlates with expression of WT1 in the smooth muscle wall as well as in the stromal cells (decidua cells). There is no WT1 expression in villous trophoblasts or extravillous trophoblasts including the endovascular trophoblasts. There appears no WT1 expression in uterine NK cells or lymphocytes (data not shown), although WT1 functions are known to be important for myeloid cells and myeloid leukemia [28].

Spiral artery remodeling in early pregnancy is critical for normal successful pregnancy, and the two separate molecular mechanisms are proposed: trophoblasts-dependent or trophoblasts-independent [3, 29]. Trophoblasts-dependent remodeling is associated with direct invasion of extravillous trophoblasts into the spiral artery with replacement of smooth muscle wall and endovascular trophoblasts plugs in the lumen [7]. WT1 gene function is not known to play any role in the process of trophoblastic invasion into the vessels, as there is no WT1 expression in trophoblasts or NK cells. It is unclear if these remodeled arteries contain the adventitia layer, but no WT1 expression was observed in the entire transformed arterial wall. However, the smooth muscle hypertrophy/hyperplasia of trophoblasts-independent spiral artery remodeling appears to correlate with WT1 gene expression in the smooth muscle cells in early pregnancy. Both of the mechanisms results in narrowing of the vascular lumen, thus limiting the maternal blood flow into the intervillous spaces and embryo, leading to hypoxic environment which was shown to be critical for fetal development [30]. Furthermore, in third trimester after the trophoblasts dependent smooth muscle remodeling, the spiral artery restores the endothelial layers and the smooth muscle layers within the decidua and superficial (inner) layers of myometrium, and the restoration of endothelium and smooth muscle cells appears associated with increased WT1 expression (Figure 5). WT1 and WT1 associated protein (WTAP) have been shown to regulate vascular smooth muscle proliferation [31, 32]. The restoration of spiral artery is also associated with regression (involution, de-plugging) of endovascular trophoblasts.

The adventitia layer of muscular arteries is equivalent to the serosal layer of visceral organs, and the serosal surface is covered by mesothelium which expresses WT1 [33]. In mice model, WT1 gene deletion resulted in fetal death due to abnormal development of vasculature [9, 19, 33]. In young adult mice WT1 deletion lead to multiple organ failures including renal failure, diminished splenic erythropoiesis, bone and fat development as well as abnormal circulating IGF1 and cytokines levels [9, 19]. The serosal mesothelial cells served as progenitor cells not only to the visceral fat tissue but also to the smooth muscle cells of the vessels such as coronary arteries, and WT1 is critical for maintenance of mesothelial progenitor cells [33–36]. Based on the current data, the endothelial cells appear important to restoration of smooth muscle wall of the spiral artery after trophoblasts-dependent remodeling, and no adventitia layer or serosal mesothelial layers were observed for spiral artery of endometrium, placental stem villous arteries or umbilical vessels. Spiral artery remodeling during pregnancy and restoration after pregnancy related to WT1 functions requires more in vitro and animal model studies, and this process is likely important for further risk of cardiovascular disease after pregnancy.

It is interesting to note that WT1 expression was reduced in adenomyosis (endometriosis). It is known that defective decidualization of the endometrium is related to endometriosis and IGF family members appear important for decidualization [37–39]. WT1 regulates expression of multiple IGFs, IGF binding proteins and IGF receptors, and the WT1 function appears more complex in physiological condition during menstrual cycles and pregnancy [19, 40]. It is also interesting to note that WT1 expression is abundant in the tubal epithelium but not in tubal stromal cells, and the fallopian tubes do not undergo decidualization during pregnancy. The fact that tubal pregnancy occurs occasionally raises an interesting question whether decidualization of endometrium is absolutely required for embryonic implantation, and how embryonic implantation occurs in tubal pregnancy in the absence of decidualization. Finally, WT1 is a gene with multiple functions and the upstream factors influencing the WT1 gene function/expression in regeneration of endothelial cells and smooth muscle cells as well as in endometriosis will likely yield important information not only about the decidual vasculopathy and preeclampsia but also pathogenesis of endometriosis.

## References

1. Benirschke, K., G.J. Burton, and R.N. Baergen Pathology of the Human Placenta. 6th ed. 2012: Springer.

2. Cunningham, F.G., Williams obstetrics. 25th edition. ed. 2018, New York: McGraw-Hill. xvi, 1328 pages.

3. Pijnenborg, R., L. Vercruysse, and M. Hanssens, The uterine spiral arteries in human pregnancy: facts and controversies. Placenta, 2006. 27(9-10): p. 939–58.

4. Harris, L.K., et al., Trophoblast-and vascular smooth muscle cell-derived MMP-12 mediates elastolysis during uterine spiral artery remodeling. Am J Pathol, 2010. 177(4): p. 2103–15.

5. Harris, L.K., et al., Placental bed research: II. Functional and immunological investigations of the placental bed. Am J Obstet Gynecol, 2019. 221(5): p. 457–469.

6. Zhang, P., Decidual vasculopathy and spiral artery remodeling revisited II: relations to trophoblastic dependent and independent vascular transformation. J Matern Fetal Neonatal Med, 2020: p. 1–7.

7. Zhang, P., Phenotypic Switch of Endovascular Trophoblasts in Decidual Vasculopathy with Implication for Preeclampsia and Other Pregnancy Complications. Fetal Pediatr Pathol, 2020: p. 1–20.

8. Craven, C.M., T. Morgan, and K. Ward, Decidual spiral artery remodelling begins before cellular interaction with cytotrophoblasts. Placenta, 1998.19(4): p. 241–52.

9. Hastie, N.D., Wilms’ tumour 1 (WT1) in development, homeostasis and disease. Development, 2017. 144(16): p. 2862–2872.

10. Call, K.M., et al., Isolation and characterization of a zinc finger polypeptide gene at the human chromosome 11 Wilms’ tumor locus. Cell, 1990. 60(3): p. 509–20.

11. Haber, D.A., et al., An internal deletion within an llpl3 zinc finger gene contributes to the development of Wilms’ tumor. Cell, 1990. 61(7): p. 1257–69.

12. Haber, D.A. and D.E. Housman, The genetics of Wilms’ tumor. Adv Cancer Res, 1992. 59: p. 41–68.

13. Haber, D.A. and A.J. Buckler, WT1: a novel tumor suppressor gene inactivated in Wilms’ tumor. New Biol, 1992. 4(2): p. 97–106.

14. van den Heuvel-Eibrink, M.M., Wilms Tumor. 2016.

15. Buckler, A.J., et al., Isolation, characterization, and expression of the murine Wilms’ tumor gene (WT1) during kidney development. Mol Cell Biol, 1991.11(3): p. 1707–12.

16. Pelletier, J., et al., Expression of the Wilms’ tumor gene WT1 in the murine urogenital system. Genes Dev, 1991. 5(8): p. 1345–56.

17. Patek, C.E., et al., A zinc finger truncation of murine WT1 results in the characteristic urogenital abnormalities of Denys-Drash syndrome. Proc Natl Acad Sci USA, 1999. 96(6): p. 2931–6.

18. Francke, U., et al., Aniridia-Wilms’ tumor association: evidence for specific deletion of 11p13. Cytogenet Cell Genet, 1979. 24(3): p. 185–92.

19. Chau, Y.Y., et al., Acute multiple organ failure in adult mice deleted for the developmental regulator Wt1. PLoS Genet, 2011. 7(12): p. elOO24O4.

20. Kaspar, H.G. and C.P. Crum, The utility of immunohistochemistry in the differential diagnosis of gynecologic disorders. Arch Pathol Lab Med, 2015. 139(1): p. 39–54.

21. Husain, A.N., et al., Guidelines for Pathologic Diagnosis of Malignant Mesothelioma 2017 Update of the Consensus Statement From the International Mesothelioma Interest Group. Arch Pathol Lab Med, 2018. 142(1): p. 89–108.

22. Khong, T.Y., et al., Sampling and Definitions of Placental Lesions: Amsterdam Placental Workshop Group Consensus Statement. Arch Pathol Lab Med, 2016. 140(7): p. 698–713.

23. Gebrane-Younes, J., N.M. Hoang, and L. Orcel, Ultrastructure of human umbilical vessels: a possible role in amniotic fluid formation? Placenta, 1986. 7(2): p. 173–85.

24. Anthony, F.W., et al., Progesterone up-regulates WT1 mRNA and protein, and alters the relative expression of WT1 transcripts in cultured endometrial stromal cells. J Soc Gynecol Investig, 2003. 10(8): p. 509–16.

25. Werner, H., et al., Transcriptional repression of the insulin-like growth factor I receptor (IGF-I-R) gene by the tumor suppressor WT1 involves binding to sequences both upstream and downstream of the IGF-I-R gene transcription start site.J Biol Chem, 1994. 269(17): p. 12577–82.

26. Roberts, C.T., Control of insulin-like growth factor (IGF) action by regulation of IGF-I receptor expression.Endocr J, 1996. 43 Suppl: p. S49–55.

27. Moffett, P., et al., Antagonism of WT1 activity by protein self-association.Proc Natl Acad Sci U S A, 1995. 92(24): p. 11105–9.

28. Pronier, E., et al., Genetic and epigenetic evolution as a contributor to WT1-mutant leukemogenesis.Blood, 2018. 132(12): p. 1265–1278.

29. Moser, G., et al., Human trophoblast invasion: new and unexpected routes and functions.Histochem Cell Biol, 2018.150(4): p. 361–370.

30. Scholz, H. and K.M. Kirschner, Oxygen-Dependent Gene Expression in Development and Cancer: Lessons Learned from the Wilms’ Tumor Gene, WT1.Front Mol Neurosci, 2011. 4: p. 4.

31. Small, T.W., et al., Wilms’ tumor 1-associating protein regulates the proliferation of vascular smooth muscle cells.Circ Res, 2006. 99(12): p. 1338–46.

32. Small, T.W., L.O. Penalva, and J.G. Pickering, Vascular biology and the sex of flies: regulation of vascular smooth muscle cell proliferation by wilms’ tumor 1-associating protein.Trends Cardiovasc Med, 2007. 17(7): p. 230–4.

33. Martínez-Estrada, O.M., et al., Wt1 is reguired for cardiovascular progenitor cell formation through transcriptional control of Snail and E-cadherin.Nat Genet, 2010. 42(1): p. 89–93.

34. Chau, Y.Y. and N.D. Hastie, The role of Wt1 in regulating mesenchyme in cancer, development, and tissue homeostasis.Trends Genet, 2012. 28(10): p. 515–24.

35. Chau, Y.Y., et al., Visceral and subcutaneous fat have different origins and evidence supports a mesothelial source.Nat Cell Biol, 2014.16(4): p. 367–75.

36. Chau, Y.Y. and N. Hastie, Wt1, the mesothelium and the origins and heterogeneity of visceral fat progenitors.Adipocyte, 2015. 4(3): p. 217–21.

37. Matsuzaki, S., et al., Expression of WT1 is down-regulated in eutopic endometrium obtained during the midsecretory phase from patients with endometriosis.Fertil Steril, 2006. 86(3): p. 554–8.

38. Sbracia, M., et al., Differential expression of IGF-I and IGF-II in eutopic and ectopic endometria of women with endometriosis and in women without endometriosis.Am J Reprod Immunol, 1997. 37(4): p. 326–9.

39. Gurgan, T., et al., Serum and peritoneal fluid levels of IGF I and II and insulinlike growth binding protein-3 in endometriosis.J Reprod Med, 1999. 44(5): p. 450–4.

40. Reeve, A.E., et al., Expression of insulin-like growth factor-II transcripts in Wilms’ tumour.Nature, 1985. 317(6034): p. 258–60.

